# Quantitative Neuropeptidomics Reveals Thermal Acclimation-Induced Remodeling of Peptidergic Signaling in the American Lobster *Homarus americanus*

**DOI:** 10.64898/2026.03.07.710231

**Authors:** Vu Ngoc Huong Tran, Sonal Kedia, Gaoyuan Lu, Kendra G. Selby, Thao Duong, Zachary Del Mundo, Eve Marder, Lingjun Li

## Abstract

Global warming and rising ocean temperatures pose substantial challenges to marine ecosystems and crustacean populations. As an ectothermic species, the American lobster (*Homarus americanus*) relies on physiological and neurochemical mechanisms to maintain homeostasis under varying environmental conditions. To elucidate the role of neuropeptides in neuronal plasticity and systemic adaptation to temperature fluctuations, we employed a quantitative mass spectrometry-based approach to probe key neuropeptides involving thermal adaptation in four lobster neural tissues at three temperatures: 4 °C (cold), 11 °C (control), and 18 °C (warm). Peptidomic profiling revealed a global reduction in peptide abundance during cold exposure, alongside coordinated, tissue-specific reconfigurations of the neuropeptidome between experimental groups. Cold exposure led to a significant downregulation of RFamide, leucokinin, and pyrokinin peptides in the commissural ganglia, whereas B-type allatostatin (AST-B), natalisin, and RYamide peptides were drastically elevated in the brain of warm-acclimated animals, with comparatively fewer detectable peptide abundance changes in the sinus gland and the stomatogastric ganglion. Collectively, our findings elucidate neuropeptide signaling pathways underlying thermal tolerance and adaptive resilience in *Homarus americanus*, offering insights into the survival mechanism and neurochemical basis of neural circuits in response to thermal acclimation.

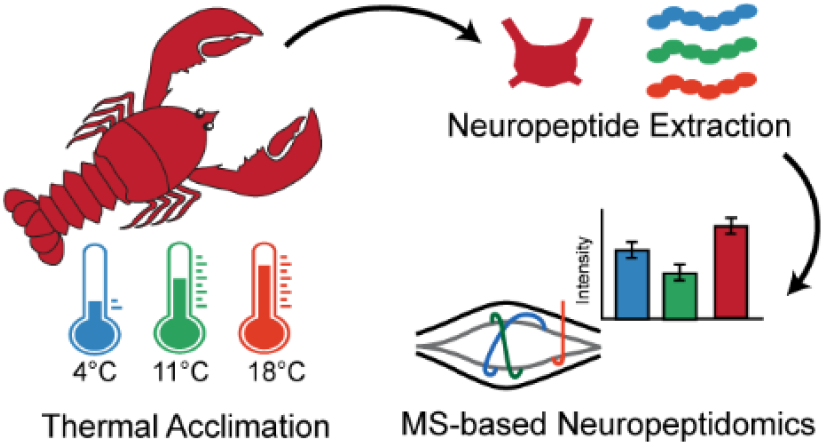

## INTRODUCTION

Climate change is one of the most urgent global challenges, with rising temperatures not only transforming ecosystems and threatening biodiversity but also having profound effects on both human and animal health. While much attention has been paid to physical consequences such as sea level rise and habitat loss, thermal stress also impacts biological systems in more subtle but critical ways, particularly by challenging neural functions across species. In humans, heat stress has been associated with worsened neurological symptoms, increased hospitalizations for dementia and stroke, and higher prevalence of mood disorders and cognitive dysfunction.^1-5^ Similar vulnerabilities are observed in animals, where thermal perturbations can impair neural development, synaptic plasticity, and sensorimotor integration, ultimately compromising behaviors essential for survival such as feeding, locomotion, and predator avoidance.^6-9^ These effects are pronounced in marine ectothermic organisms, whose body temperature closely follows their surrounding conditions.^10, 11^ Since temperature directly governs biochemical reaction rates and influences cellular signaling, even modest temperature shifts can profoundly disrupt neuronal excitability, synaptic transmission, and circuit stability, resulting in destabilized network activity and altered behavioral output.^12-14^ Altogether, temperature is a key regulator of neural signaling and stability across phyla, and it is important to understand how neural circuits adapt to or fail under prolonged thermal stress.

The American lobster, *Homarus americanus*, offers a powerful model for probing the influence of temperature on neural systems. As a marine ectotherm, the lobster is highly sensitive to ambient temperature, making it ecologically relevant for studying neural stress under climate change. Numerous studies have shown temperature influences on the lobster physiological parameters (*e*.*g*., glucose, protein, and metabolite concentration), developmental stages, food consumption, and the outputs of lobster neuronal circuits, including the pyloric rhythm resilience and cardiac performance.^15-24^ Most importantly, lobsters possess a diverse and evolutionarily conserved neuropeptide repertoire, including peptides homologous to those in mammals, that has been extensively characterized using mass spectrometry (MS), providing a strong foundation for elucidating temperature-dependent neuromodulatory functions.^25-28^ Neuropeptides are the key signaling interface between environmental stimuli and neural functions, with broad roles in regulating many physiological processes such as pain, stress, appetite, and environmental influences.^29-36^ By modulating ion channels, synaptic strength, and network dynamics, neuropeptides contribute critically to circuit stability and adaptability to varying internal and external conditions.^37^

MS-based neuropeptidomics enables simultaneous detection and quantification of endogenous neuropeptides from small tissue samples and circulating fluids, capturing both known and previously uncharacterized peptides and allowing temperature-dependent shifts in neuropeptide expression to be assessed at a systems level.^38, 39^ Substantial progress has been made in understanding how temperature affects neural function, particularly in crustaceans. A previous neuropeptidomics study in the Jonah crab, *Cancer borealis*, demonstrated that acute temperature elevation led to significant decreases in RFamide, RYamide, and orcokinin neuropeptide abundance and discovered a temperature-stress marker peptide in the hemolymph, indicating neuropeptide involvement in thermal stress responses.^40^ However, this work focused on short-term challenges to extremely high temperature rather than long-term acclimation to a wider range of temperatures in the ocean. In parallel, peptidomics study in hibernating ground squirrels revealed large, season and hibernation stage-dependent shifts in hypothalamic and pituitary neuropeptides, demonstrating that temperature-linked physiological states are accompanied by extensive neuropeptide remodeling in mammals.^41^ However, whether comparable neuropeptidomic adaptations occur in invertebrates, and how they contribute to circuit-level thermal robustness remains largely unexplored and is well suited for investigation in the simple, well-defined neural circuits of crustaceans.

In this study, an MS-based peptidomics approach was employed to characterize the dynamic changes in the neuropeptidome landscape of the American lobster nervous system in response to prolonged thermal acclimation. By acclimating animals to cold and warm temperature extremes, we identified neuropeptidomic signatures associated with adaptive, compensatory, and behavioral responses to temperature perturbation. Collectively, this work provides a framework for understanding how chronic exposure to extreme temperatures reshapes neuromodulatory signaling to support neural adaptability and resilience at the molecular level, offering insights into the survival mechanism and neurochemical basis of neural circuits in response to thermal stress.

## MATERIALS AND METHODS

### Chemicals and Materials

All chemicals and solvents, unless noted otherwise, were purchased from Thermo Fisher Scientific (Pittsburgh, PA) or MilliporeSigma (St. Louis, MO).

### Animals and Tissue Collection

American lobsters, *Homarus americanus*, were purchased from Commercial Lobster (Boston, MA) and housed for at least three weeks prior to any experiments. The animal tanks were made with Instant Ocean Sea Salt and maintained at 11°C with alternating 12h light/12h dark cycles. A total of eighteen lobsters were acclimated for three weeks at 18 °C (*N* = 7; warm group), 11 °C (*N* = 5; control group), or 4 °C (*N* = 6; cold group). Prior to dissection, all animals were placed on ice for 30 minutes to induce anesthesia. Four distinct neural tissues, including a pair of sinus glands (SG), the brain, a pair of commissural ganglia (CoG), and the stomatogastric ganglion (STG) were collected in chilled physiological saline solution (479 mM NaCl; 12.74 mM KCl; 20 mM MgSO_4_, 3.91 mM Na_2_SO_4_, 13.67 mM CaCl_2_, 5 mM HEPES, pH 7.45). Tissues were immediately transferred into 200 μL of ice-cold acidified methanol (90:9:1, v/v/v; methanol:water:glacial acetic acid), snap-frozen on dry ice, and stored at -80°C until further analysis.

### Neuropeptide Extraction

Tissues were homogenized using a Fisher Scientific sonic dismembrator (Pittsburgh, PA) with 8-sec on/15-sec off pulses at 50% amplitude, for 7 to 12 cycles until no visible tissue chunks remained. Following homogenization, samples were centrifuged at 20,000 g at 4°C for 40 min. The supernatant was collected, dried down, subsequently reconstituted in 15 μL of water containing 0.1% formic acid (FA), and mixed with 15 μL of an internal standard consisting of a 500 pg/μL isotope-encoded orcomyotropin (FDAFTTGF[^13^C9:^15^N]GHN) solution. Samples were desalted with C_18_ Ziptips (Millipore Sigma, St. Louis, MO) according to manufacturer instructions, and peptides were sequentially eluted in 20 μL of 50% CAN in 0.1% FA, followed by 20 μL of 75% ACN in 0.1% FA. The eluates were combined and evaporated using a SpeedVac concentrator on medium heat and stored at −80°C until liquid chromatography tandem mass spectrometry (LC-MS/MS) analysis.

### LC-MS/MS Data Acquisition

Dried samples were reconstituted in 12 μL of water containing 0.1% FA prior to MS analysis. Samples were loaded onto a self-packed microcapillary column (approximately 18 cm in length, 75 μm i.d.), packed with bridged ethylene hybrid (BEH) C18 particles (1.7 μm, 130 Å, Waters). Spectra were acquired using An Orbitrap Fusion Lumos Tribrid mass spectrometer coupled to a Dionex UltiMate 3000 UPLC system (Thermo Fisher Scientific). The LC separation was performed using 0.1% FA in water as mobile phase A and 0.1% FA in 80% acetonitrile as mobile phase B, operating at a constant flow rate of 0.3 μL/min. Peptide separation was achieved using a 60-min gradient as follows: 0-18 min 3% B, 18-40 min 18-45% B, 40-45 min 45-99% B, 45-55 min 99% B, 55-55.5 min 99 - 3% B, 55.5-60 min 3% B. MS data were acquired from 300−1500 *m/z* at a resolving power of 60k, normalized automatic gain control (AGC) target of 50%, and maximum injection time of 100 ms. Data were acquired in data-dependent acquisition (DDA) mode, with the 20 most intense precursors being selected for MS^2^ fragmentation with a normalized collision energy (NCE) of 30%, a resolving power of 15k, a normalized AGC target of 20%, and a maximum injection time of 100 ms. Precursors were subject to dynamic exclusion for 45 s with a 10-ppm tolerance.

### Database Search and Data Analysis

The MS raw data were searched in PEAKS Studio 13 (Bioinformatics Solutions Inc.) against a meticulously curated database containing known neuropeptide precursors filtered from the genome-derived *Homarus americanus* proteome database downloaded from Uniprot on Feb 12^th^, 2023. The search parameters include 10 ppm parent mass error tolerance, 0.02 Da fragment mass error tolerance, and an unspecific digest mode. A maximum of three variable PTMs including methionine oxidation, N-terminal pyro-glutamate formation from glutamic acid and glutamine residues, and C-terminal amidation were allowed. The identified peptides were filtered using the following thresholds: Proteins -10lgP ≥ 0 and ≥1 unique peptide; Peptides/PSM -10lgP ≥ 15, with -10lgP referring to a PEAKS confidence score. The label free quantification (LFQ) was conducted using the PEAKS Q module with the following criteria: 10 ppm precursor mass error tolerance, peptide quality score ≥ 5, detection in a minimum of two samples per experimental group, and normalization based on internal standard signal ratios. Perseus (version 1.6.0.7) was used for log2(x) transformation and handling missing values by imputation from a normal distribution. Data processing and visualization were further conducted using Microsoft Excel and custom Python scripts. For statistical comparison, MS-detected peak areas were evaluated between the acclimation and control (11 °C) groups using Welch’s t-test with unequal variances. Peptides were considered significantly altered if they exhibited an absolute log_2_ fold change smaller than 0.5 and a *p*-value below 0.05. One brain and SG samples originating from a lobster in the control group were excluded from all analyses due to failure during the sample collection stage.

## RESULTS AND DISCUSSION

The natural habitat of *Homarus americanus* is characterized by wide seasonal temperature fluctuations spanning from near-freezing conditions to approximately 25 °C, influenced by oceanographic factors such as seasonal cycles, wind patterns, and tidal movements.^42^ A behavioral thermoregulation study demonstrated that American lobsters preferentially occupy water temperatures of approximately 12.5 °C in the laboratory condition and avoid temperatures above 19 °C, consistent with the moderate conditions they experience in their natural coastal habitats.^43^ Therefore, this study employed lobsters acclimated to an control temperature of 11 °C, and those acclimated to more extreme temperatures of 4 °C and 18 °C were designated as cold and warm groups, respectively. It is noteworthy that we did not conduct formal behavioral or morphological assessments of lobsters acclimated to different temperatures to reduce additional handling or disturbance that could influence experimental conditions. However, informal observations during routine animal care indicated clear temperature-associated differences in activity and appearance, which were taken into account in downstream data interpretations. Specifically, warm-acclimated lobsters appeared more active and aggressive, whereas cold-acclimated lobsters were comparatively hypoactive compared to the control group. Differences in carapace coloration were also noted, with warm-acclimated animals appearing brighter red than their cold-acclimated counterparts. Following three weeks of acclimation, four lobster neural (brain, CoG, and STG) and neuroendocrine (SG) tissues were collected, extracted for neuropeptides, and subjected to MS analysis (**Figure 1**). These tissues were selected for their critical involvement in neuropeptide synthesis, secretion, and modulation of neural circuit activity.

**Figure 1.**
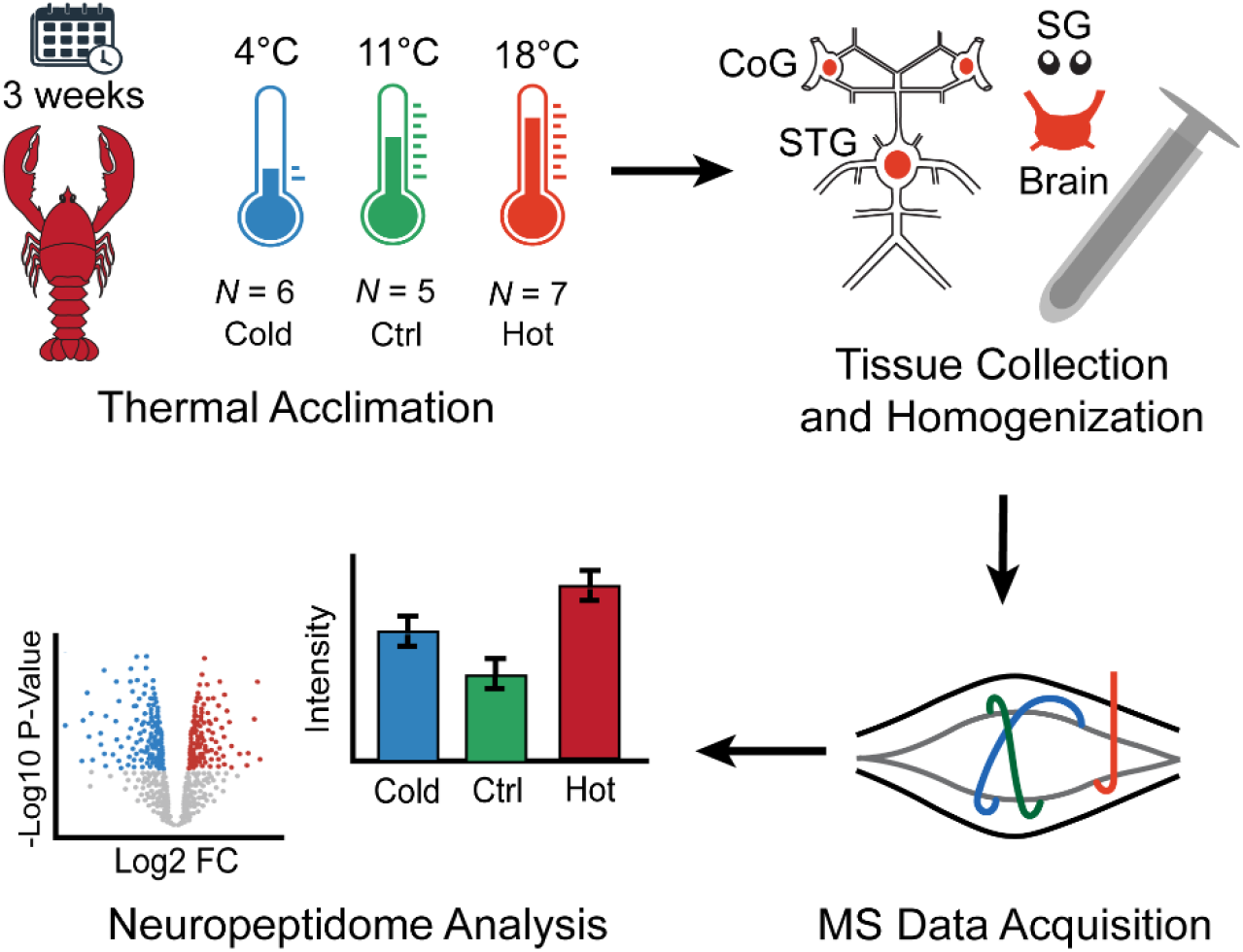
Scheme of the experimental workflow. American lobsters (*Homarus americanus*) were acclimated to three temperature conditions (18°C, 11°C, or 4°C). Four neural tissues were collected from each animal and processed for mass spectrometry-based neuropeptidomics analysis. CoG: commissural ganglia, STG: stomatogastric ganglion, SG: sinus gland.

### Peptidome Identification and Features

In our MS-based neuropeptidomics workflow, raw MS spectra were searched against a curated *H. americanus* genome-derived database comprising 88 carefully filtered neuropeptide prohormones. Using permissive identification parameters consistent with PEAKS software guidelines to enable comprehensive peptide detection, our study detected a total of 1,834 peptides in the SG, followed by 1,026 in the brain, 911 in the CoG, and 316 in the STG (**Figure 2A, Data File S1**). To enhance quantitative reliability, more stringent criteria were applied for quantification, yielding approximately 100-300 confidently quantifiable peptides per tissue (**Figure 2A, Data File S2**). Quantifiable peptides were defined as those reliably measured in the MS^1^ spectra in at least two biological replicates per experimental group and possessed a quality score greater than 5. We further examined the family distribution of these quantifiable peptides across all four selected tissues (**Figure 2B**). Most peptides were derived from well-recognized crustacean neuropeptide families such as orcokinin, crustacean hyperglycemic hormone B (CHH-B), and A- and B-type allatostatin (AST-A, AST-B). Their prevalence across tissues suggests their central roles in the peptidergic signaling system of the American lobster, consistent with previous neuropeptidomics profiling studies.^26, 27^ We next investigated the tissue distribution of the quantified peptides, identifying twelve peptides present in all four selected tissues (**Figure 2C**). Most of these represent mature neuropeptides, defined as those carry the signature sequence motif of their family and follow the canonical processing pattern involving cleavage at mono- or di-basic residues (K/R)^44^, including six AST-A peptides, the well-characterized Val^1^-SIFamide, two orcokinins, and one RYamide. Despite this shared core set, a substantial proportion of peptides displayed tissue-restricted expression patterns, supporting functional specialization of neuropeptidergic signaling across different neural and neuroendocrine tissues. Lastly, a key advantage of MS-based neuropeptidomics is the capacity to directly identify post-translational modifications (PTMs), which are critical determinants of neuropeptide bioactivity. In this study, we detected numerous peptides bearing the most prevalent neuropeptide PTMs, including C-terminal amidation and N-terminal pyroglutamylation of glutamine or glutamic acid residues (**Figure 2D**). Notably, eleven peptides carried both modifications simultaneously, forming so-called “capped” peptides. Such dual-terminal processing is a hallmark of mature, bioactive neuropeptides, as these modifications enhance resistance to proteolytic degradation, stabilize peptide conformation, and are often required for high-affinity receptor interactions.^45^ The prevalence of capped peptides and other modified peptides supports the biological relevance of the detected peptides and confirms that our workflow effectively captured functionally active neuropeptide forms rather than degradation products or biosynthetic intermediates.

**Figure 2.**
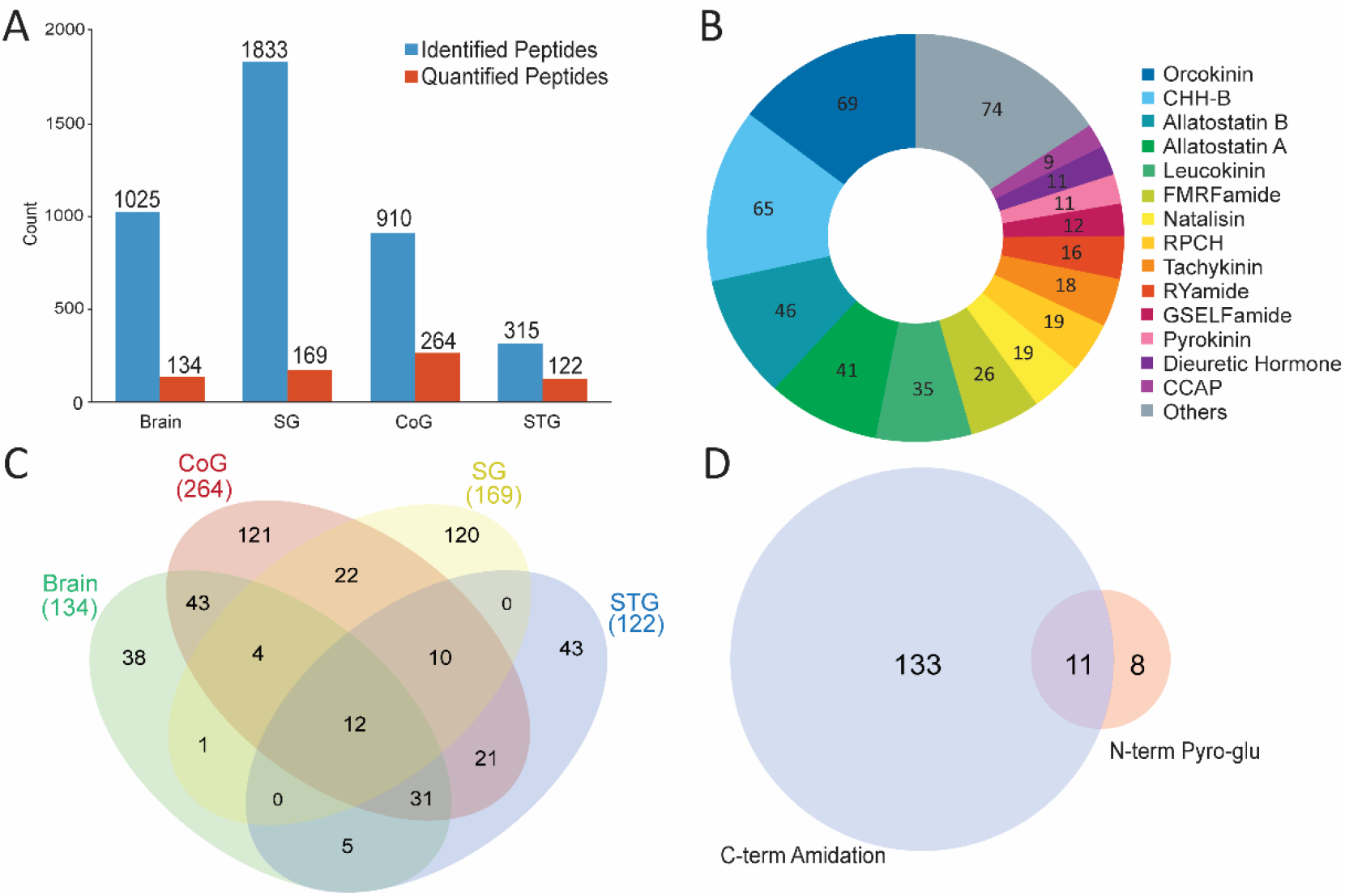
Global neuropeptidome features across four lobster neural tissues. (A) Bar graph displaying the number of identified and quantified peptides in four neural tissues. (B) The family distribution and (C) Overlap of quantified peptides across all four selected tissues. (D) Venn diagram showing the overlap between quantified peptides carrying the C-terminal amidation or N-terminal pyro-glutamylation. Abbreviations: sinus glands (SG), commissural ganglia (CoG), stomatogastric ganglion (STG), crustacean hyperglycemic hormone B (CHH-B), red pigment concentrating hormone (RPCH), crustacean cardioactive peptide (CCAP).

### Overall Thermal Acclimation-Induced Peptidomic Changes

We next performed quantitative analysis of the confidently quantified peptides to probe global and tissue-specific neuropeptidomic remodeling in response to thermal acclimation. To assess the global changes in peptide levels, volcano plot analyses comparing warm *versus* cold (**Figure 3A**), warm *versus* control (**Figure 3B**), and cold *versus* control (**Figure 3C**) were conducted. These comparisons revealed that the neural peptidome undergoes a profound and highly structured reconfiguration in response to temperature changes. The most substantial peptidomic divergence was observed along the thermal gradient, characterized by a dominant reduction in peptide abundance in the cold group relative to the warmer conditions. This observation suggests that cold acclimation may trigger strategic peptidomic silencing, potentially serving as a metabolic conservation mechanism to reduce the energetic costs of neuropeptide biosynthesis and secretion under low temperature conditions. Meanwhile, the warm *versus* control comparison revealed a slightly more stable peptidome with fewer changes in peptide abundance among the four selected tissues, indicating that the transition from 11 °C to 18 °C remains within a robust homeostatic range which requires only targeted modulation rather than a global peptidomic reconfiguration. Notably, a distinct level of thermal sensitivity was observed among the four neural tissues. The CoG exhibited the highest number of significantly altered peptides across all temperature transitions (**Figure 3**, red), suggesting that it acts as a primary neuroendocrine integrator that senses thermal input to coordinate systemic responses. The brain (**Figure 3**, green) displayed an intermediate variability with a regulatory trend similar to the CoG, where a significant proportion of peptides was upregulated in the warm group relative to their cold counterparts, raising the possibility that the brain and CoG may share conserved modulatory mechanisms for driving thermal compensation. In contrast, the SG (**Figure 3**, yellow) and STG (**Figure 3**, blue) showed comparatively stable peptidomic profiles, with only a small subset of peptides showing downregulation at 18 °C relative to those at 4 °C. Note that the relatively small tissue mass and overall lower peptide abundance in SG and STG could reduce the sensitivity in label-free MS quantification and limit the detection of subtle changes. Overall, these observable patterns support a model in which temperature adaptation involves coordinated modulation of peptide signaling to rebalance circuit excitability and metabolic state under varying thermal conditions.

**Figure 3.**
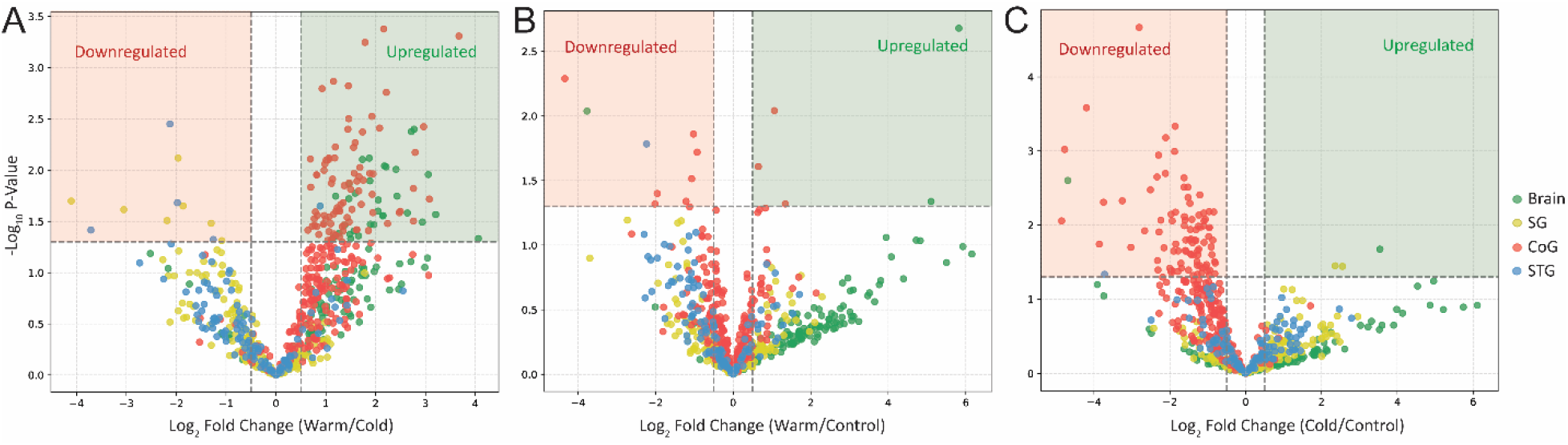
Comparative volcano plots of significantly changed peptides across thermal acclimation states. Pairwise comparisons are shown for (A) warm *versus* cold, (B) warm *versus* control, and (C) cold *versus* control groups across four neural tissues. The y-axis represents the -log10(*p*-value), and the x-axis represents the log2(Fold Change) in peptide abundance. Horizontal dashed lines indicate the significance threshold (*p* < 0.05), while vertical dashed lines denote the fold-change cut-offs (absolute log2FC > 0.5). Shaded areas highlight significantly downregulated (red) and upregulated (green) peptides. Abbreviations: SG: sinus glands; CoG: commissural ganglia; STG: stomatogastric ganglion.

### Tissue-specific Neuromodulatory Responses to Thermal Adaptation

We visualized the log_2_ fold change analysis of selected significantly altered neuropeptides in each tissue to reveal distinct tissue-specific neuroendocrine and neuromodulatory response profiles. The SG, which is one of the principal neuropeptide hormone release sites in the lobster, exhibited selective thermal modulation of neuroendocrine peptides (**Figure 4A, Table S1**). Specifically, a sulfakinin peptide (GGGEYDDYGHLRFamide) was significantly reduced in warm-acclimated lobsters relative to cold conditions, suggesting enhanced secretion into the hemolymph at elevated temperature. This unsulfated sequence most likely corresponds to Hoa-SK (II) whose mature form contains a tyrosine sulfation, which is notoriously labile during MS fragmentation, leading to a predominant detection of the desulfated form. Sulfakinins are homologous to vertebrate cholecystokinin (CCK)-like peptides and regulate feeding, gut motility, locomotor activity, and aggression in arthropods.^25, 46^ The reduced Hoa-SK (II) abundance in the SG warm-acclimated lobsters therefore suggests increased endocrine signaling that may contribute to the heightened behavioral activity observed during elevated thermal stress. A similar trend of Hoa-SK (II) abundance alterations were also observed in the CoG (**Figure 4D**) but not in the brain (**Figure 4C**) or STG (**Figure 4B**) across the thermal gradients. In crustaceans, crustacean hyperglycemic hormones (CHHs) are released from the sinus gland (SG) into the hemolymph during environmental stressors such as elevated temperature, acute hypoxia, or metal exposure, typically accompanied by increased glucose levels in the hemolymph.^47-49^ Consistently, our data revealed that two CHH-B fragment peptides were significantly reduced in the SG of warm-acclimated lobsters compared to cold-acclimated animals. This reduction may reflect enhanced release of CHH peptides from the SG under prolonged exposure to higher temperatures to regulate hyperglycemia and osmoregulatory responses. Moreover, while two red pigment concentrating hormone (RPCH) precursor-related peptides exhibited a significant decrease in the SG of warm animals, the well-studied mature RPCH peptide itself (pQLNFSPGWamide, indicated by the red star in **Figure 4A**) followed a similar downward trend that did not reach statistical significance (*p* = 0.13). This discrepancy may stem from complex precursor processing dynamics or the inherent technical challengesof reliably quantifying subtle abundance shifts in such a small neuroendocrine tissue analyzed by label-free MS approach. While RPCH is primarily known for its role as a chromatophorotropin regulating pigment aggregation, a previous study demonstrated that it can also indirectly induce hyperglycemia in crustaceans, likely through stimulation of CHH release from the SG.^50^ Our data, which show a concurrent decrease in both RPCH and CHH peptides in the SG of warm-acclimated animals, may therefore reflect a coordinated neuroendocrine response under the chronic exposure to higher temperatures. Additionally, an orcokinin was also observed with significant down regulation in the warm-acclimated lobsters relative to their cold-acclimated counterparts. This pattern alligns with previous observations in the Jonah crab (*Cancer borealis*) SG which found that under acute temperature elevation, several orcokinin peptides also showed reduced abundance, further corroborating that orcokinin signaling may be involved in thermal stress responses.^40^

**Figure 4.**
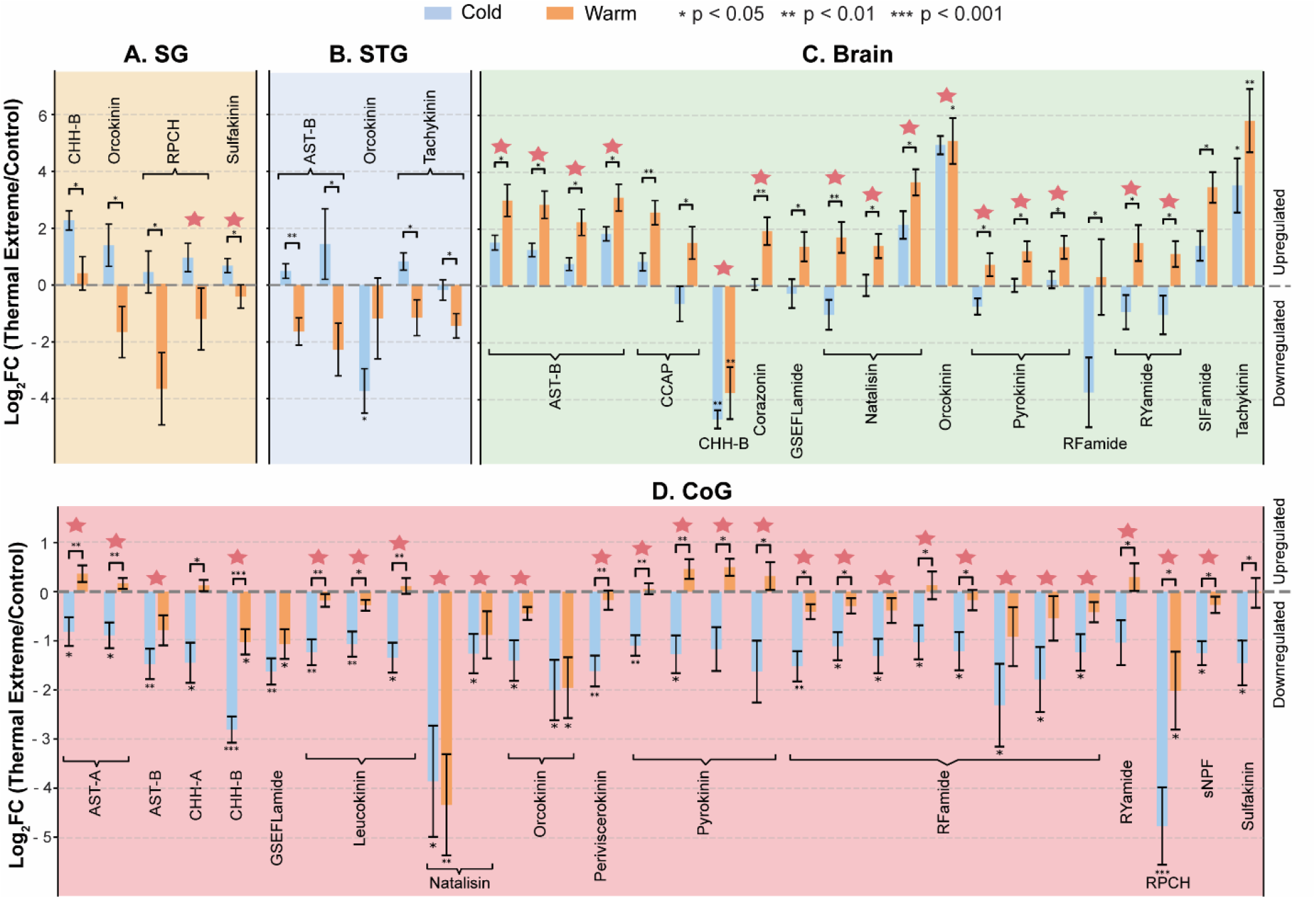
Bar charts showing the log_2_ fold changes in peptide abundance relative to the control temperature (11 °C) for cold-acclimated (4 °C) and warm-acclimated (18 °C) groups in (A) sinus glands (SG), (B) stomatogastric ganglion (STG), (C) Brain, and (D) commissural ganglia (CoG). Positive values indicate increased abundance relative to control, whereas negative values indicate decreased abundance. Error bars represent the standard error of the mean (SEM) for the thermal-acclimated group. Statistical significance between thermal and control groups was assessed using Welch’s t-test. Asterisks above individual bars denote significant differences relative to control. Brackets indicate significant differences between cold and warm groups. For both statistical analyses, * indicates p < 0.05, ** indicates p < 0.01, and *** indicates p < 0.001. Red stars denote mature neuropeptides that carry the signature sequence motif of the family and follow the canonical processing pattern involving cleavage at dibasic residues (K/R), consistent with typical neuropeptide prohormone maturation. Abbreviations: AST-A: A-tyoe allatostatin; AST-B: B-type Allatostatin; CHH-A: Crustacean hyperglycemic hormone A; CHH-B: Crustacean hyperglycemic hormone B; CCAP: Crustacean cardioactive peptide; RPCH: Red pigment-concentrating hormone; sNPF: short Neuropeptide F.

On the other hand, the STG revealed several key neuropeptide families that undergo significant temperature-dependent shifts in abundance profiles (**Figure 4B, Table S2**). A truncated orcokinin (1–12) peptide (NFDEIDRSGFGF) was significantly downregulated in the cold-acclimated STG relative to controls. In *H. americanus*, this detected fragment represents the conserved N-terminal core shared by three 13-residue *H. americanus* orcokinin variants, which differ only at the last C-terminal residue.^27^ A prior electrophysiological work demonstrated that exogenous [Ala^13^]-orcokinin halves the lateral pyloric neuron spike/burst output in *H. americanus* STG, indicating a suppressive effect on pyloric rhythm.^51^ The decrease in endogenous orcokinins under cold conditions indicates that this neuromodulatory peptide family may be linked to the STG’s response to cold acclimation. Furthermore, a truncated C-terminal tachykinin fragment (PSGFLGM(O)Ramide), representing the conserved bioactive motif of two tachykinin isoforms APSGFLGMRamide and TPSGFLGMRamide, was also significantly reduced in the warm group relative to the cold. Tachykinins have been reported to activate the pyloric rhythm in other crustacean species,^52, 53^ suggesting that temperature-dependent changes in tachykinin abundance may reflect modulation of tachykinin signaling in the STG under different thermal conditions. Furthermore, AST-B neuropeptides, known as myoinhibitory peptides (MIPs), are known inhibitors of STG activity in the Jonah crab.^54^ Nonetheless, the bioactive mature AST-Bs did not show any changes in abundance, whereas some AST-B precursor related peptides (PRPs) reduced significantly in the warm-acclimated animals. This suggests PRPs and mature neuropeptides of this neuromodulatory family may be differentially regulated, indicating complex prohormone cleavage and peptidergic control in maintaining circuit stability under thermal stress. Overall, temperature-dependent shifts in neuropeptide abundance in the STG act to rebalance excitation and inhibition, supporting stability of pyloric circuit output across thermal conditions.

Thermal acclimation responses in the brain were characterized by a coordinated upregulation of diverse neuropeptide families in warm-acclimated animals, suggesting a global shift in neuroendocrine state (**Figure 4C, Table S3**). Notably, mature neuropeptides derived from the same precursors (*e*.*g*., AST-B, RYamide, pyrokinin, and natalisin) exhibited synchronized increases in abundance in the warm *versus* cold lobsters. This suggests that temperature-driven regulation occurs at the level of prohormone synthesis or processing, ensuring the co-release of multiple bioactive peptides. Furthermore, while the bioactive crustacean cardioactive peptide (CCAP) (PF<u>C</u>NAFTG<u>C</u>amide; underlined cysteines denote intramolecular disulfide bond) contains a disulfide bond that hinders direct mass spectrometric detection, its associated peptides derived from the same precursor showed a significant upregulation in the warm animals, implying a concomitant rise in CCAP levels in the brain during warm acclimation. Additionally, corazonin was significantly upregulated in the 18 °C group (*p* < 0.01), identifying it as a key neuroendocrine responder to high-temperature acclimation. Since its functional roles in crustaceans remain largely uncharacterized, the pronounced thermal sensitivity observed here warrants further investigation into its physiological targets as a component of the crustacean thermal stress response. The brain (**Figure 4C**) also exhibited a significant thermal response in RYamide signaling, with two mature isoforms showing increased abundance in warm-acclimated animals relative to the cold group. Notably, a similar trend regarding RYamide regulation was obsered in the CoG (**Figure 4D**), which may indicate concerted signaling efforts by these two tissues. In a previous study on the effect of acute temperature elevation on the Jonah crab, RYamides were downregulated in their primary neuroendocrine organs (*i*.*e*., pericardial organs) in the acute thermal stress group.^40^ The opposing changes observed in this study suggest tissue-specific regulation of RYamide signaling, showing distinct neuroendocrine and central neuromodulatory roles during thermal adaptation. Lastly, warmer temperatures also serve as a critical environmental factor for seasonal mating and molting cycles in lobsters.^55^ The observed enrichment of natalisins in the brains of the warm-acclimated group provides the first indirect functional evidence of this newly discovered neuropeptide’s role at higher temperatures, likely representing reproductive readiness and mating signaling as demonstrated in other arthropods.^56^ Collectively, these findings indicate that the brain undergoes a systematic neuropeptide processing and reconfiguration to sustain neural performance, locomotor activity, and reproduction regulation as the animal approaches its upper thermal limits.

The CoG emerged in this study as the most responsive neural tissue to thermal acclimation, with 129 significantly altered neuropeptides representing the largest degree of peptidomic reconfiguration among all four examined tissues (**Figure 4D, Table S4**). The CoG is a key component of the stomatogastric nervous system (STNS) and contains modulatory projection neurons that provide descending input to the STG, contributing to the generation and modulation of pyloric and gastric mill motor patterns.^57^ In this study, many neuropeptides detected in the CoG belong to families whose members have been shown to regulate the pyloric rhythms, feeding-related behaviors, or metabolic processes in crustaceans.^58-61^ Several mature members of these families exhibited significant decreases in the CoG under cold acclimation, including eight RFamide peptides, three leucokinins, and one short neuropeptide F. While RFamide peptides were also detected in the STG, their abundance remained unchanged in that tissue. This distribution pattern suggests that temperature-dependent regulation of these peptides may occur primarily at the level of upstream modulatory centers of the STNS. Additionally, four mature pyrokinins and a recently identified periviscerokinin^27^ were also significantly reduced in the CoG during cold acclimation, highlighting the broad peptidergic remodeling occurring within this upstream modulatory ganglion and demonstrating the first evidence linking these peptide families to the crustacean thermal response that warrants future investigations into their roles in environmental adaptation. Given that descending inputs from the CoG are critical for STG dynamics, these localized shifts in peptide abundance in the CoG likely reflect a remodeling of the modulatory drive available to the motor circuit under different thermal conditions.

## CONCLUSION

This study comprehensively characterized the dynamics of the neuropeptidomic landscape in *Homarus americanus* under prolonged thermal acclimation. Using label-free high-resolution MS, we successfully profiled hundreds of endogenous peptides and quantified their temperature-dependent shifts across the central nervous system. Notably, we observed a significant downregulation of multiple RFamide, leucokinin, and pyrokinin peptides in the CoG during cold exposure, while AST-B, natalisin, and RYamide levels were markedly elevated in the brain of warm-acclimated animals. In contrast, fewer peptidomic changes were detected in the SG and STG, potentially reflecting the inherent technical challenges of quantifying subtle abundance shifts in these smaller tissues using a label-free approach. These abundance shifts likely stem from a complex interplay between peptide synthesis, axonal transport, and release kinetics to facilitate the systemic metabolic demands and balance the local circuit stability under temperature perturbation. These peptidomic signatures corroborate previous literature regarding the established roles of these neuropeptide families in regulating reproduction, growth, and homeostasis. Ultimately, the results revealed a highly coordinated, tissue-specific reconfiguration of the peptidergic signaling under temperature perturbation and underscored the role of neuropeptides as the molecular thermostats, providing fundamental principles for understanding how environmental fluctuations shape neural circuit resilience, physiological adaptation, and homeostatic regulation in more complex mammalian systems.

## Data Availability

The mass spectrometry proteomics data have been deposited to the ProteomeXchange Consortium via the MassIVE partner repository with dataset identifier **MSV000101008**.

## Author Contributions

Conceptualization: V.N.H.T., S.K., E.M. and L.L.; Experimentation: V.N.H.T., S.K., G.L., K.S. and T.D.; Data analysis and presentation: V.N.H.T.; Writing (Original draft): V.N.H.T., Z.D.M., E.M. and L.L.; Funding acquisition: E.M. and L.L. All the authors read and commented on the manuscript.

## Notes

The authors declare no competing financial interest.

## Acknowledgements

This work was supported in part by the National Science Foundation (CHE-2108223) and the National Institutes of Health (NIH) through Grants R01DK071801, R01NS029346, and R35NS097343. The Orbitrap instruments were purchased through the support of a NIH shared instrument grant (S10RR029531) and the Office of the Vice Chancellor for Research and Graduate Education at the University of Wisconsin-Madison. Z.D.M. was supported in part by the NIH Biotechnology Training Grant (T32 GM135066) through the University of Wisconsin-Madison. L.L. would like to acknowledge NIH grants R01AG078794, R01AG052324, S10OD028473, and S10OD025084, the Research Forward grant by University of Wisconsin - Madison Office of the Vice Chancellor for Research with funding from the Wisconsin Alumni Research Foundation, as well as funding support from a Vilas Distinguished Achievement Professorship and Charles Melbourne Johnson Professorship with funding provided by the Wisconsin Alumni Research Foundation and University of Wisconsin-Madison School of Pharmacy.

