## Supplemental Information for "Quantitative Neuropeptidomics Reveals Thermal Acclimation-Induced Remodeling of Peptidergic Signaling in the American Lobster *Homarus americanus*"

---

[*] Corresponding author

 (L. L)

**Table of Contents**

**Table S1.** List of significantly changed peptides during thermal acclimation in the *H. americanus* SG.

**Table S2.** List of significantly changed peptides during thermal acclimation in the *H. americanus* STG.

**Table S3.** List of significantly changed peptides during thermal acclimation in the *H. americanus* brain.

**Table S4.** List of significantly changed peptides during thermal acclimation in the *H. americanus* CoG.

**Table S1.** List of significantly changed peptides during thermal acclimation in the *H. americanus* SG. Statistical significance between groups was assessed using Welch's t-test. The significance threshold is set at $p < 0.05$. Mature neuropeptides are defined as those that carry the signature sequence motif of the family and follow the canonical processing pattern involving cleavage at dibasic residues (K/R), consistent with typical neuropeptide prohormone maturation. AST-A: Allatostatin A; CHH-B: Crustacean hyperglycemic hormone B; RPCH: Red pigment-concentrating hormone.

| **Family** | **Peptide** | **Mature** | ***p*-Value** | **$Log_2FC$** | **Comparison** |
|---|---|---|---|---|---|
| AST-A | EVSDDDHDEDEQDIGVEEEMS | No | 0.031 | -2.184 | Warm/Cold |
| CHH-B | VEGVSRMEKLLSSISPSSTPLGFLSQDHSVN | No | 0.022 | -1.863 | Warm/Cold |
| CHH-B | LSSISPSSTPLGFLSQDHS | No | 0.033 | -1.298 | Warm/Cold |
| Orcokinin | PIKAAPARSSPQQDAAA | No | 0.035 | 2.359 | Cold/Control |
| Orcokinin | GPIKAAPARSSPQQDAAAGYTDGAPamide | No | 0.036 | 2.552 | Cold/Control |
| Orcokinin | PIKAAPARSSPQQDAAAGYTDGAPV | Yes | 0.024 | -3.051 | Warm/Cold |
| RPCH | AAAASGTDPAAAS | No | 0.008 | -1.962 | Warm/Cold |
| RPCH | AAAASGTDPAAASLHPAPPAVLTAASGAN | No | 0.020 | -4.104 | Warm/Cold |
| Sulfakinin | GGGEYDDYGHLRFamide | Yes | 0.049 | -1.086 | Warm/Cold |

**Table S2.** List of significantly changed peptides during thermal acclimation in the *H. americanus* STG. Statistical significance between groups was assessed using Welch's t-test. The significance threshold is set at $p < 0.05$. Mature neuropeptides are defined as those that carry the signature sequence motif of the family and follow the canonical processing pattern involving cleavage at dibasic residues (K/R), consistent with typical neuropeptide prohormone maturation. AST-B: Allatostatin B; CHH-B: Crustacean hyperglycemic hormone B

| **Family** | **Peptide** | **Mature** | ***p*-Value** | **$Log_2FC$** | **Comparison** |
|---|---|---|---|---|---|
| AST-B | PHLEDAQLDAA | No | 0.004 | -2.126 | Warm/Cold |
| AST-B | SSSSSPQQDDPASSPSHIEE | No | 0.038 | -3.711 | Warm/Cold |
| CHH-B | RSVEGVSRM(+15.99)EKLLSSISPSSTPLGFLSQDHSVN | Yes | 0.022 | 0.885 | Warm/Cold |
| Orcokinin | NFDEIDRSGFGF | No | 0.046 | -3.720 | Cold/Control |
| Orcokinin | GPIKAAPARSSPQQ | No | 0.016 | -2.231 | Warm/Control |
| Tachykinin | GEGQDTPQDRE | No | 0.021 | -1.977 | Warm/Cold |
| Tachykinin | PSGFLGM(O)Ramide | No | 0.047 | -1.257 | Warm/Cold |

**Table S3.** List of significantly changed peptides during thermal acclimation in the *H. americanus* brain. Statistical significance between groups was assessed using Welch's t-test. The significance threshold is set at $p < 0.05$. Mature neuropeptides are defined as those that carry the signature sequence motif of the family and follow the canonical processing pattern involving cleavage at dibasic residues (K/R), consistent with typical neuropeptide prohormone maturation. AST-A: Allatostatin A; AST-B: Allatostatin B; CHH-B: Crustacean hyperglycemic hormone B; CCAP: Crustacean cardioactive peptide.

| Family | Peptide | Mature | *p*-Value | Log$_2$FC | Comparison |
|---|---|---|---|---|---|
| AST-A | ESDEDSDKRSQMYSFGLamide | No | 0.009 | 2.209 | Warm/Cold |
| AST-B | PDNTRVSPR | No | 0.010 | 2.408 | Warm/Cold |
| AST-B | APDMMSVAAPNQ | No | 0.011 | 3.055 | Warm/Cold |
| AST-B | APDMMSVAAPNQA | No | 0.013 | 1.884 | Warm/Cold |
| AST-B | NNWRSLQGSWamide | Yes | 0.019 | 1.464 | Warm/Cold |
| AST-B | GWTLWGKRPDNTRVSPR | No | 0.027 | 3.204 | Warm/Cold |
| AST-B | PHLEDAQLDAAEV | No | 0.039 | 1.566 | Warm/Cold |
| AST-B | SADWNKLRGAWamide | Yes | 0.041 | 1.271 | Warm/Cold |
| AST-B | VGWSSMHGTWamide | Yes | 0.042 | 1.482 | Warm/Cold |
| AST-B | AWNKLQGAWamide | Yes | 0.017 | 1.589 | Warm/Cold |
| CCAP | DIGDLLEGKD | No | 0.008 | 1.736 | Warm/Cold |
| CCAP | STPHTQPRQHLTSTPQQKVETEKQ | No | 0.028 | 2.147 | Warm/Cold |
| CHH-B | RSVEGVSRMEKLLSSISPSSTPLGFLSQDHSVN | Yes | 0.002 | -4.691 | Cold/Control |
| | | | 0.009 | -3.768 | Warm/Control |
| Corazonin | pQTFQYSRGWTNamide | Yes | 0.008 | 1.871 | Warm/Cold |
| GSEFLamide | MAMSNPAGRYKSPQLLNRGV | No | 0.034 | 1.964 | Warm/Cold |
| GSEFLamide | ALGSEFLGKRVMGSEFLamide | No | 0.044 | 1.654 | Warm/Cold |
| Leucokinin | SDPLLPASQHEPNT | No | 0.026 | 2.637 | Warm/Cold |
| Natalisin | SFWAMRamide | Yes | 0.004 | 2.717 | Warm/Cold |
| Natalisin | EGEETHPFWVS | No | 0.034 | 1.893 | Warm/Cold |
| Natalisin | EGEAPPFWVSRamide | Yes | 0.034 | 1.383 | Warm/Cold |
| Natalisin | EGEETHPFWVSRamide | Yes | 0.047 | 1.499 | Warm/Cold |
| Neuropeptide F | KYLQELamide | No | 0.025 | 2.048 | Warm/Cold |
| Orcokinin | GPIKAAPARSSPQQDAAAGYTDGAPV | Yes | 0.046 | 5.104 | Warm/Control |
| Pyrokinin | SDFAFSPRLamide | Yes | 0.014 | 1.468 | Warm/Cold |
| Pyrokinin | ADFAFSPRLamide | Yes | 0.018 | 1.197 | Warm/Cold |
| Pyrokinin | GDGFAFSPRLamide | Yes | 0.046 | 1.148 | Warm/Cold |
| RFamide | LLKYFLPASQAWGGDAYPIGQEGT | No | 0.047 | 4.059 | Warm/Cold |
| RYamide | SSPSQGLPEI | No | 0.004 | 2.771 | Warm/Cold |
| Ryamide | pQGFYTQRYamide | Yes | 0.018 | 2.429 | Warm/Cold |
| RYamide | FIGGSRYamide | Yes | 0.028 | 2.144 | Warm/Cold |
| SIFamide | VYRKPPFNGSIF | No | 0.020 | 2.052 | Warm/Cold |
| Tachykinin | APSGFLGM(O)RG | No | 0.032 | 2.937 | Warm/Cold |
| Tachykinin | APSGFLGMRG | No | 0.009 | 2.177 | Warm/Cold |
| Tachykinin | AGEGQDTPQDRE | No | 0.021 | 3.541 | Cold/Control |
| | | | 0.002 | 5.821 | Warm/Control |

**Table S4.** List of significantly changed peptides during thermal acclimation in the *H. americanus* CoG. Statistical significance between groups was assessed using Welch's t-test. The significance threshold is set at $p < 0.05$. Mature neuropeptides are defined as those that carry the signature sequence motif of the family and follow the canonical processing pattern involving cleavage at dibasic residues (K/R), consistent with typical neuropeptide prohormone maturation. ALP: Agatoxin-like peptide; AST-A: Allatostatin A; AST-B: Allatostatin B; CHH-A: Crustacean hyperglycemic hormone A; CHH-B: Crustacean hyperglycemic hormone B; CCAP: Crustacean cardioactive peptide; DH: diuretic hormone; RPCH: Red pigment-concentrating hormone; sNPF: short Neuropeptide F.

| Family | Peptide | Mature | *p*-Value | $Log_2FC$ | Comparison |
|---|---|---|---|---|---|
| ALP | DDVAGSDPIK | No | 0.033 | -1.710 | Cold/Control |
| AST-A | AKYSFGIamide | No | 0.008 | 1.007 | Warm/Cold |
| AST-A | ADPYAFGLGKKAGQYSFGLamide | No | 0.007 | -0.843 | Cold/Control |
| | | | 0.015 | 0.698 | Warm/Cold |
| AST-A | AGHYAFGLamide | Yes | 0.028 | 0.832 | Warm/Cold |
| AST-A | AGRYAFGLamide | Yes | 0.031 | -0.821 | Cold/Control |
| AST-A | DPDM(O)DM(O)DKRPRDYAFGLamide | No | 0.009 | 1.064 | Warm/Control |
| AST-A | DPDM(O)DMDKRPRDYAFGLamide | No | 0.042 | 1.160 | Warm/Cold |
| AST-A | DPDMDMDKRPRDYAFGLamide | No | 0.007 | -1.851 | Cold/Control |
| AST-A | DVSITEDTLE | No | 0.003 | 1.926 | Warm/Cold |
| AST-A | DVSITEDTLED | No | 0.045 | 1.272 | Warm/Cold |
| AST-A | EGLYSLGLD | No | 0.0003 | -4.199 | Cold/Control |
| | | | 0.0005 | 3.671 | Warm/Cold |
| AST-A | ESDEDSDKRSQMYSFGLamide | No | 0.0005 | -1.860 | Cold/Control |
| | | | 0.002 | 1.453 | Warm/Cold |
| AST-A | HSNYGFGLamide | Yes | 0.045 | -0.814 | Cold/Control |
| | | | 0.008 | 1.179 | Warm/Cold |
| AST-A | KVEDGASPRT | No | 0.030 | -2.327 | Cold/Control |
| AST-A | PRNYAFGLamide | Yes | 0.024 | -0.890 | Cold/Control |
| | | | 0.008 | 1.058 | Warm/Cold |
| AST-A | SDLYSFGLGKKSGSYNFGLamide | No | 0.007 | -1.105 | Cold/Control |
| | | | 0.006 | 1.190 | Warm/Cold |
| AST-A | SDSDSDQYTLamide | Yes | 0.048 | -1.383 | Cold/Control |
| | | | 0.032 | 1.469 | Warm/Cold |
| AST-A | SKLYGFGLG | No | 0.048 | 1.346 | Warm/Control |
| AST-A | SVGDLPEVSKVEDGASPRT | No | 0.014 | 1.371 | Warm/Cold |
| AST-A | VSKVEDGASPRT | No | 0.011 | 1.882 | Warm/Cold |
| AST-B | APDMM(+15.99)SVAAPNQ | No | 0.047 | -0.961 | Cold/Control |
| AST-B | DDPASSPSHIEE | No | 0.038 | 1.623 | Warm/Cold |
| AST-B | GEELQAAED | No | 0.026 | 2.474 | Warm/Cold |
| AST-B | PHLEDAQL | No | 0.023 | -1.571 | Cold/Control |
| AST-B | PHLEDAQLDA | No | 0.043 | 1.199 | Warm/Cold |
| AST-B | PHLEDAQLDAAE | No | 0.009 | -1.937 | Cold/Control |
| AST-B | PHLEDAQLDAAEV | No | 0.017 | -1.097 | Cold/Control |

| | | | | | |
|---|---|---|---|---|---|
| AST-B | SSSSPQQDDPASSPSHIE | No | 0.040 | -1.508 | Cold/Control |
| AST-B | TQVSSSSSSPQQDDPASSPSHIEE | No | 0.012 | -2.663 | Cold/Control |
| | | | 0.025 | 2.494 | Warm/Cold |
| AST-B | APDMMSVAAPN | No | 0.025 | -1.436 | Cold/Control |
| AST-B | APDMMSVAAPNQ | No | 0.016 | -1.581 | Cold/Control |
| | | | 0.023 | 1.231 | Warm/Cold |
| AST-B | ASDWGQFRamide | Yes | 0.022 | 1.711 | Warm/Cold |
| AST-B | APDMMSVAAPNQA | No | 0.031 | -1.282 | Cold/Control |
| | | | 0.022 | 1.302 | Warm/Cold |
| AST-B | DMMSVAAPNQA | No | 0.016 | 1.660 | Warm/Cold |
| AST-B | DWNKLRGAWamide | Yes | 0.005 | -1.470 | Cold/Control |
| AST-B | PDNTRVSPR | No | 0.009 | 1.683 | Warm/Cold |
| AST-B | NNWRSLQGSWamide | Yes | 0.041 | -0.949 | Cold/Control |
| AST-C | KALPDQDPQVY | No | 0.033 | 2.240 | Warm/Cold |
| CCAP | DIGDLLEGKD | No | 0.028 | 0.970 | Warm/Cold |
| CCAP | LTSTPQQKVETE | No | 0.005 | -3.254 | Cold/Control |
| | | | 0.004 | 2.963 | Warm/Cold |
| CCAP | LTSTPQQKVETEKQ | No | 0.009 | -2.237 | Cold/Control |
| | | | 0.017 | 1.930 | Warm/Cold |
| CCAP | pQHLTSTPQQKVETEKQ | No | 0.020 | -3.028 | Cold/Control |
| | | | 0.019 | 3.077 | Warm/Cold |
| CCAP | STPHTQPRQHLTSTPQQKVETE | No | 0.005 | -3.749 | Cold/Control |
| | | | 0.015 | 2.757 | Warm/Cold |
| CCAP | STPHTQPRQHLTSTPQQKVETEKQ | No | 0.009 | -4.854 | Cold/Control |
| CCAP | TSTPQQKVETEKQ | No | 0.022 | -1.895 | Cold/Control |
| | | | 0.004 | 2.079 | Warm/Cold |
| CHH-A | RSVEGASRM(+15.99)EKL | No | 0.042 | 0.859 | Warm/Cold |
| CHH-A | RSVEGASRM(O)EKLLSSSNSPSSTPLGFLSQDHSV | No | 0.015 | -1.446 | Cold/Control |
| | | | 0.011 | 1.570 | Warm/Cold |
| CHH-A | RSVEGASRM(O)EKLLSSSNSPSSTPLGFLSQDHSVN | Yes | 0.030 | 0.768 | Warm/Cold |
| CHH-B | RSVEGVSRMEKLLSSISPSSTPLGFLSQDHSVN | Yes | 0.00002 | -2.809 | Cold/Control |
| | | | 0.001 | 1.787 | Warm/Cold |
| | | | 0.014 | -1.022 | Warm/Control |
| CHH-B | RSVEGVSRM(+15.99)EKLLSSISPSSTPLGFLSQDHSVN | Yes | 0.002 | 0.926 | Warm/Cold |
| CHH-B | RSVEGVSRM(+15.99)EKLLSSISPSSTPLGFLSQDHSV | No | 0.035 | 0.778 | Warm/Cold |
| CCRFamide | VSDLQSELGPDYDDSTamide | No | 0.010 | -1.618 | Cold/Control |
| | | | 0.0004 | 2.162 | Warm/Cold |
| DH | SSDDGLDLHHDDNLYAQDQAADLAESS | No | 0.002 | -2.117 | Cold/Control |
| | | | 0.034 | 1.366 | Warm/Cold |
| RFamide | APSKNFLRFamide | Yes | 0.034 | 0.961 | Warm/Cold |
| RFamide | APVPPVVAALDPPTDALLPAQSQEDDLFALPE | No | 0.008 | 0.692 | Warm/Cold |
| | | | 0.007 | -0.959 | Cold/Control |
| RFamide | DGSDDYPSSSSSAESPAPVVVVRPVEYPRYV | No | 0.044 | -1.163 | Cold/Control |
| | | | 0.021 | 1.351 | Warm/Cold |

| | | | | | |
|---|---|---|---|---|---|
| RFamide | DQNRNFLRFamide | Yes | 0.014 | -0.976 | Cold/Control |
| RFamide | FSHDRNFLRFamide | Yes | 0.042 | -1.787 | Cold/Control |
| RFamide | GAHKNYLRFamide | Yes | 0.040 | -2.312 | Cold/Control |
| RFamide | GDRNFLRFamide | Yes | 0.035 | -1.211 | Cold/Control |
| | | | 0.049 | 1.038 | Warm/Cold |
| RFamide | GNRNFLRFamide | Yes | 0.012 | -1.310 | Cold/Control |
| RFamide | GYSDRNYLR | No | 0.038 | -1.693 | Cold/Control |
| RFamide | GYSDRNYLRFamide | Yes | 0.010 | -1.112 | Cold/Control |
| | | | 0.036 | 0.823 | Warm/Cold |
| RFamide | HAAPVPPVVAALDPPTDALLPAQSQEDDLFALPE | No | 0.016 | -0.866 | Cold/Control |
| RFamide | LLKYFLPASQAWGGDAYPIGQEGT | No | 0.039 | 0.641 | Warm/Cold |
| RFamide | PSKNFLRFamide | Yes | 0.034 | -1.031 | Cold/Control |
| | | | 0.027 | 1.158 | Warm/Cold |
| RFamide | SDRNYLRFamide | Yes | 0.004 | -1.520 | Cold/Control |
| | | | 0.013 | 1.113 | Warm/Cold |
| RFamide | SDTNDYEGEEM(O)PESPE | No | 0.011 | -1.460 | Cold/Control |
| RFamide | SDTNDYEGEEMPESPE | No | 0.007 | -1.965 | Cold/Control |
| | | | 0.013 | 1.752 | Warm/Cold |
| RFamide | SGSPMEFATDLQEDVELPVEE | No | 0.003 | -1.526 | Cold/Control |
| | | | 0.019 | -0.927 | Warm/Control |
| RFamide | NRNFLRFamide | Yes | 0.021 | -1.237 | Cold/Control |
| GSEFLamide | AALPTHLPDELDDPVV | No | 0.035 | 0.729 | Warm/Cold |
| GSEFLamide | ALGSEFLGKRVMGSEFLamide | No | 0.003 | -1.626 | Cold/Control |
| | | | 0.031 | -1.068 | Warm/Control |
| GSEFLamide | pQYEPEFAHTLDYDT | No | 0.012 | 1.134 | Warm/Cold |
| Orcokinin | AAGPIKAAPARSSPQQDAAAGYTDGAPV | No | 0.030 | 0.999 | Warm/Cold |
| Orcokinin | GPIKAAP | No | 0.042 | -1.970 | Cold/Control |
| Orcokinin | GPIKAAPA | No | 0.044 | -1.955 | Cold/Control |
| Orcokinin | GPIKAAPARSSPQQ | No | 0.022 | -1.781 | Cold/Control |
| Orcokinin | GPIKAAPARSSPQQD | No | 0.033 | 1.101 | Warm/Cold |
| Orcokinin | GPIKAAPARSSPQQDA | No | 0.024 | 1.525 | Warm/Cold |
| Orcokinin | GPIKAAPARSSPQQDAAAamide | No | 0.022 | -1.591 | Cold/Control |
| | | | 0.039 | 1.346 | Warm/Cold |
| Orcokinin | GPIKAAPARSSPQQDAAAGYTDGAPV | Yes | 0.017 | -1.403 | Cold/Control |
| Orcokinin | GPIKAAPARSSPQQDAAAGYTDGAPVK | No | 0.038 | -2.003 | Cold/Control |
| | | | 0.040 | -1.955 | Warm/Control |
| Sulfakinin | EFDEYGHMRFamide | Yes | 0.004 | -1.210 | Cold/Control |
| | | | 0.025 | 1.280 | Warm/Cold |
| Sulfakinin | GGGEYDDYGHLRFamide | Yes | 0.024 | -1.452 | Cold/Control |
| | | | 0.029 | 1.428 | Warm/Cold |
| Leucokinin | AEGTLSRLSESTLKAALDENSPEDNVDN | No | 0.041 | -1.008 | Cold/Control |
| Leucokinin | AEGTLSRLSESTLKAALDENSPEDNVDNH | No | 0.008 | -1.419 | Cold/Control |
| Leucokinin | ASPISEDSQLSDLYTSQ | No | 0.002 | -1.625 | Cold/Control |
| Leucokinin | LPINDWGN | No | 0.012 | -2.283 | Cold/Control |

| | | | | | |
|---|---|---|---|---|---|
| Leucokinin | LPINDWGNKRVPFSTWGamide | No | 0.001 | -1.876 | Cold/Control |
| Leucokinin | QAFHPWGamide | Yes | 0.016 | -1.344 | Cold/Control |
| | | | 0.003 | 1.460 | Warm/Cold |
| Leucokinin | RTFSAWAamide | Yes | 0.011 | -0.956 | Cold/Control |
| | | | 0.011 | 0.817 | Warm/Cold |
| Leucokinin | SDIDEKRPSFNAWAamide | Yes | 0.011 | -1.192 | Cold/Control |
| | | | 0.006 | 1.555 | Warm/Cold |
| Leucokinin | SDPLLPASQHEPN | No | 0.016 | -1.314 | Cold/Control |
| | | | 0.029 | 1.067 | Warm/Cold |
| Leucokinin | SDPLLPASQHEPNT | No | 0.036 | -1.065 | Cold/Control |
| Leucokinin | SDSDEKRPSFSAWAamide | Yes | 0.022 | -1.266 | Cold/Control |
| | | | 0.025 | 1.204 | Warm/Cold |
| Leucokinin | SESNEKRPSFNAWAamide | No | 0.003 | -2.512 | Cold/Control |
| Leucokinin | SPSM(O)DLSGNQ | No | 0.046 | -1.274 | Cold/Control |
| | | | 0.013 | 1.470 | Warm/Cold |
| Leucokinin | SPSM(O)DLSGNQD | No | 0.040 | 1.062 | Warm/Cold |
| Leucokinin | SPSMDLSGNQ | No | 0.013 | 1.624 | Warm/Cold |
| Leucokinin | SPSMDLSGNQD | No | 0.009 | -1.602 | Cold/Control |
| | | | 0.024 | 1.266 | Warm/Cold |
| Leucokinin | SSDVNLQDGEDDEPVSAWIGRRLQ | No | 0.015 | 1.498 | Warm/Cold |
| Leucokinin | SSDVNLQDGEDDEPVSAWIGRRLQDANTDD | No | 0.001 | -2.298 | Cold/Control |
| | | | 0.005 | 1.589 | Warm/Cold |
| Leucokinin | SSETDKRQGFSAWAamide | No | 0.001 | -2.109 | Cold/Control |
| | | | 0.046 | -1.214 | Warm/Control |
| Leucokinin | SSGDELDDHFLD | No | 0.048 | 1.123 | Warm/Cold |
| Leucokinin | TFRAWAamide | Yes | 0.007 | -1.234 | Cold/Control |
| | | | 0.008 | 1.056 | Warm/Cold |
| Leucokinin | TRFSAWAamide | Yes | 0.011 | -0.956 | Cold/Control |
| | | | 0.011 | 0.817 | Warm/Cold |
| Leucokinin | TRFSPWAamide | Yes | 0.008 | -1.067 | Cold/Control |
| | | | 0.026 | 0.792 | Warm/Cold |
| Periviserokinin | pQDLIPFPRVamide | Yes | 0.008 | -1.616 | Cold/Control |
| | | | 0.004 | 1.444 | Warm/Cold |
| RYamide | SDTGEVTVRSGFYANRNamide | Yes | 0.025 | -1.366 | Cold/Control |
| RYamide | SRFIGGSRYamide | Yes | 0.036 | 1.333 | Warm/Cold |
| RYamide | SSPSQGLPEI | No | 0.047 | 1.123 | Warm/Cold |
| RYamide | SSPSQGLPEIKIRSS | No | 0.020 | -0.910 | Cold/Control |
| | | | 0.034 | 0.951 | Warm/Cold |
| RYamide | SSPSQGLPEIKIRS | No | 0.002 | -2.338 | Cold/Control |
| | | | 0.011 | 1.948 | Warm/Cold |
| RYamide | SSRFIGGSRYG | No | 0.016 | -1.483 | Cold/Control |
| | | | 0.002 | 2.215 | Warm/Cold |
| Natalisin | DAVDGRAPFWISRamide | Yes | 0.018 | -3.861 | Cold/Control |
| | | | 0.005 | -4.340 | Warm/Control |

| | | | | | |
|---|---|---|---|---|---|
| Natalisin | PSSELLHQHHQ | No | 0.020 | -2.155 | Cold/Control |
| | | | 0.039 | 1.854 | Warm/Cold |
| Natalisin | QETEGNGGPFWIARamide | Yes | 0.025 | -1.261 | Cold/Control |
| Natalisin | SFWAM(O)Ramide | Yes | 0.025 | 0.651 | Warm/Control |
| Pyrokinin | AYFSPRLamide | Yes | 0.009 | -1.096 | Cold/Control |
| | | | 0.001 | 1.155 | Warm/Cold |
| Pyrokinin | GADFAFSPRLamide | Yes | 0.012 | 1.666 | Warm/Cold |
| Pyrokinin | LYYSQRPamide | Yes | 0.035 | -1.276 | Cold/Control |
| | | | 0.004 | 1.736 | Warm/Cold |
| Pyrokinin | RSEFVFSSRPamide | Yes | 0.018 | -1.845 | Cold/Control |
| | | | 0.038 | 1.618 | Warm/Cold |
| Pyrokinin | SLFSPRLamide | Yes | 0.026 | 1.945 | Warm/Cold |
| RPCH | pQLNFSPGWamide | Yes | 0.001 | -4.774 | Cold/Control |
| | | | 0.031 | 2.759 | Warm/Cold |
| | | | 0.048 | -2.015 | Warm/Control |
| sNPF | DQPAKIPTM(O)GLamide | Yes | 0.038 | -1.149 | Cold/Control |
| sNPF | DQPAKIPTMGLamide | Yes | 0.009 | 0.982 | Warm/Cold |
| | | | 0.010 | -1.250 | Cold/Control |
| SIFamide | VYRKPPFNGSIFG | No | 0.041 | -1.946 | Cold/Control |
| | | | 0.007 | 2.791 | Warm/Cold |
| SIFamide | VYRKPPFNGSIFamide | Yes | 0.033 | -0.800 | Cold/Control |
